## Supplementary figures and images for "Evaluation of nanopore sequencing for increasing accessibility of eDNA studies in biodiverse countries"

### S4 Figure

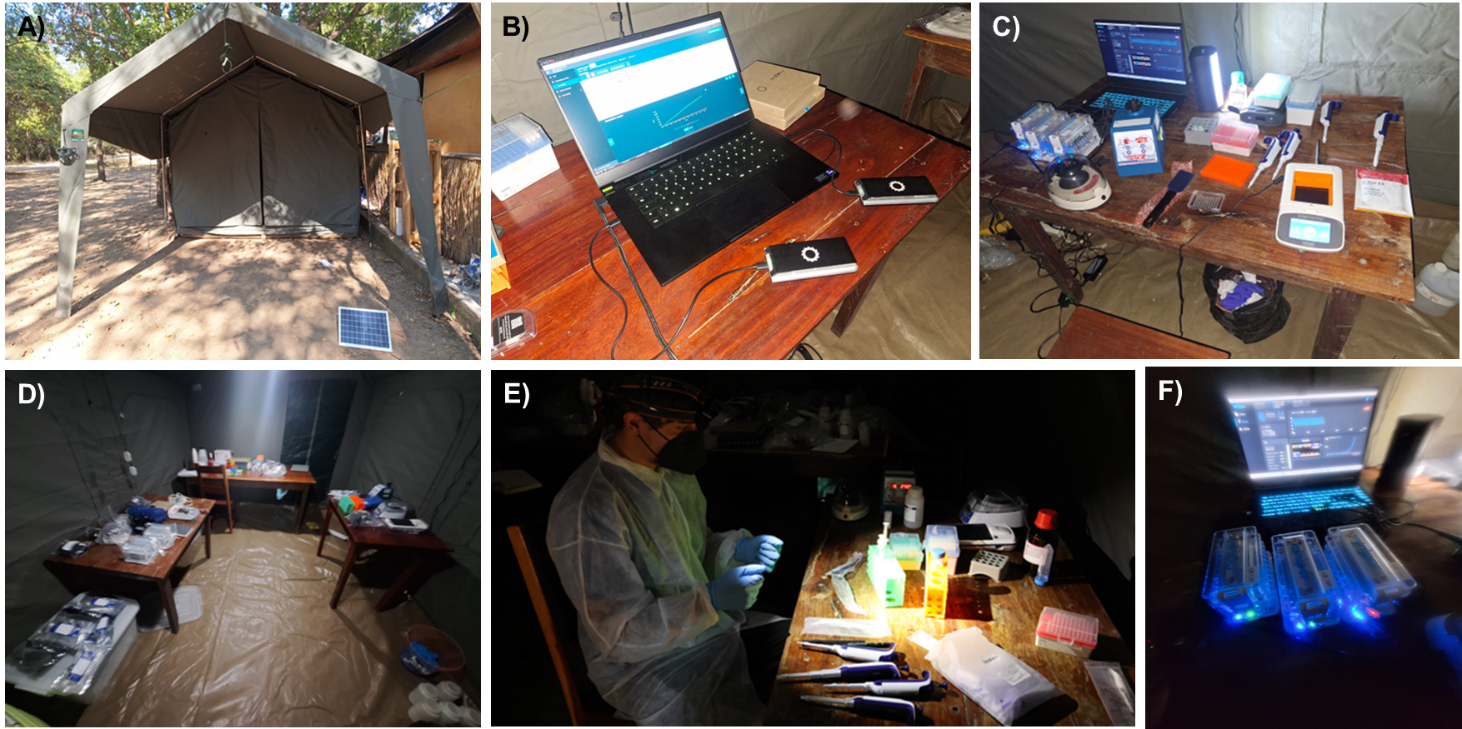
